# Genomic Polymorphisms in Toxin-Antitoxin Systems and Identification of Novel Phylo-SNPs and Polymorphisms Associated with Drug Resistance/Susceptibility in Clinical Isolates of *Mycobacterium tuberculosis* from Mumbai, India

**DOI:** 10.1101/797274

**Authors:** Kayzad Nilgiriwala, Vidushi Chitalia, Sanchi Shah, Akshata Papewar

## Abstract

Toxin-antitoxin (TA) modules are one of the prominent determinants that triggers a persistent state aiding *Mycobacterium tuberculosis* evasion to host generated stresses. The 79 characterized and putative TA systems described in *M. tuberculosis* are dominated by the VapBC, MazEF, HigAB, RelBE and ParDE TA families, largely involved in persistence and cell arrest. Hence, there is a need to maintain and conserve the TA loci in the chromosome of the pathogen. It is essential to study the genomic differences of the TA systems in clinical isolates along with its association to drug susceptibility patterns and lineage. In the current study, the TA loci and their promoter sequences were analysed from the whole genome sequence data of 74 clinical isolates. *Mykrobe Predictor* was used for lineage identification and drug resistance predictions in the clinical isolates. Polymorphisms associated with 79.8% (63/79) TA systems were observed across 72 clinical isolates. Among the TA systems, the isolates had a varying number of polymorphisms localised primarily in the toxin genes (58.7%), antitoxin genes (40.7%) and chaperones (0.6%), due to Single Nucleotide Polymorphism (SNP) resulting in transition (67.3%), transversion or frameshift mutations. Our analysis suggests the presence of novel Phylo-SNPs by establishing high confidence association of specific lineages to polymorphisms in the TA systems. Notably, association of polymorphisms in Rv1838c-1839c (VapBC13), Rv3358-3357 (YefM/YoeB) and Rv0240-0239 (VapBC24) to Delhi/Central Asia lineage. The polymorphic loci of the 3 TA systems is localised in the antitoxin gene of the Delhi/Central Asia strains, with a resultant silent mutation. The assessment of correlation between TA polymorphisms and the drug resistance profile revealed correlation of SNPs in VapBC35 with drug resistant *M. tuberculosis* strains and SNPs in VapBC24, VapBC13 and YefM/YoeB to drug sensitive strains.

## 1. INTRODUCTION

Tuberculosis (TB), a disease caused by *Mycobacterium tuberculosis*, with an annual estimate of 10.4 million new cases globally, imposes a major burden of mortality and morbidity on the human population [1]. The long duration of treatment and a large reservoir of people with latent infection are major obstacles in controlling spread of the disease. Furthermore, multidrug resistance is a deterrent for effective treatment of TB. Besides acquired drug resistance, *M. tuberculosis* exhibits modified virulence, transmissibility and pathogenicity [2]. The organism may employ endogenous mechanisms of stress evasion due to persistence of the bacilli in the infected individual. The ‘persister’ state is a dormant state wherein the bacterial cells are recalcitrant to the unfavourable conditions. The persistent cells revert back to its proliferative growth on removal of the environmental stress factor such as nutrient starvation or antibiotic presence. The persister cells are therefore not antibiotic resistant mutants, but are highly antibiotic tolerant cells [3]. An important determinant of persistence is the presence of multiple toxin-antitoxin (TA) systems [4].

The TA systems are ubiquitous in prokaryotes; and typically consist of a stable protein toxin and a relatively unstable protein or non-coding RNA antitoxin against the cognate toxin protein neutralizing the toxin. On the basis of the nature of the antitoxin and its mode of interaction, toxin-antitoxin systems have been classified into five types [5]. The TA systems in numerous organisms are associated to post-segregational killing of daughter cells devoid of plasmid in plasmid encoded TA systems; however, chromosomally encoded TA systems as in case of *M. tuberculosis*, the role of TA system seems to be central in bacterial persistence [6]. *M. tuberculosis* has the highest number of TA systems than any other known bacteria (79 characterized and putative TA systems). The involvement of TA systems in persistence is proposed to be linked to increased pathogenesis. A majority of the TA systems in *M. tuberculosis* belong to class II type that includes VapBC, MazEF, HigAB, RelBE and ParDE families of TA systems [6].

The present study analyzes the differences in the genomic structure of the TA systems of clinical isolates in comparison to H_37_Rv and correlate the polymorphisms in TA system to the drug resistance pattern and strain lineage.

## 2. MATERIALS AND METHODS

### 2.1. *M. tuberculosis* strains

A total of seventy-four clinical isolates of *M. tuberculosis* from TB patients (49 new, 12 follow-ups and 13 re-treatment cases) were used in the study [7]. Along with the clinical isolates, standard strain H_37_Rv (ATCC 27294) was also processed in triplicate.

### 2.2. Genomic DNA extraction, whole genome sequencing and analysis

Genomic DNA was isolated from the clinical strains of *M. tuberculosis* using FastPrep24 lysis method (MP Biomedicals, California, USA) as per standard protocol [7]. The extracted DNA was quantified using Qubit (Life Technologies, Carlsbad, California, USA). The DNA from the samples was processed using MiSeq Reagent Kit V2 in a MiSeq sequencer (Illumina Inc., San Diego, California, USA) as per manufacturer’s protocol producing 151 base-pair paired end reads. The sequencing data can be accessed online (NCBI, SRA Accession ID: SRP101835). Sequencing reads were assembled using *M. tuberculosis* H_37_Rv reference genome (GenBank version ID: NC_000962.3) using *Geneious* software (v10.1.3) [8], with default parameters. To diminish the possibility of assembly errors and resultant false SNPs, chimeric and low-quality reads were discarded. The sequences were aligned, visualized and analysed for toxin-antitoxin systems along with the associated SNPs, insertions and deletions which were determined with minimum coverage of 10x, Q-score of ≥ 20 and variant frequency ≥ 90%.

The TA gene loci were identified from Tuberculist database (v 2.6) and as reported by Sala *et al* [6]. Regions of 100 bp upstream for each of the 79 distinct TA gene modules was selected as the promoter sequence and analysed for polymorphisms. The polymorphisms in the TA gene modules and promoters were extracted using *R* (v 3.4.2). The table (Table 1) describes the putative or characterized TA system considered for the study. The lineage identification and drug susceptibility pattern were predicted using *Mykrobe Predictor* (v 0.4.2) [9, 10].

**Table 1.** The TA systems included in the present study.

| **TA**<br>**system ID** | **TA system** | **TA**<br>**system ID** | **TA system** |
| --- | --- | --- | --- |
| **1** | Rv0059-0060 (DarTG) | **41** | Rv1962c-1963c (VapBC35) |
| **2** | Rv0065-0064A (VapBC1) | **42** | Rv1982c-1982A (VapBC36) |
| **3** | Rv0240-0239 (VapBC24) | **43** | Rv1989c-1990c |
| **4** | Rv0277c-0277A (VapBC25) | **44** | Rv1991c-1991A (MazEF6) |
| **5** | Rv0299-0298 (MazEF10) | **45** | Rv2010-2009 (VapBC15) |
| **6** | Rv0301-0300 (VapBC2) | **46** | Rv2019-2018 |
| **7** | Rv0456A-0456B (MazEF1) | **47** | Rv2022c-2021c (HigBA2) |
| **8** | Rv0549c-0550c (VapBC3) | **48** | Rv2035-2034 |
| **9** | Rv0582-0581 (VapBC26) | **49** | Rv2063A-2063 (MazEF7) |
| **10** | Rv0595c-0596c (VapBC4) | **50** | Rv2103c-2104c (VapBC37) |
| **11** | Rv0598c-0599c (VapBC27) | **51** | Rv2142c-2142A (ParDE2) |
| **12** | Rv0609-0608 (VapBC28) | **52** | Rv2231A-2231B (vapBC16) |
| **13** | Rv0617-0616A (VapBC29) | **53** | Rv2274c-2274A (MazEF8) |
| **14** | Rv0624-0623 (VapBC30) | **54** | Rv2494-2493 (VapBC38) |
| **15** | Rv0627-0626 (VapBC5) | **55** | Rv2527-2526 (VapBC17) |
| **16** | Rv0656c-0657c (VapBC6) | **56** | Rv2530c-2530A (VapBC39) |
| **17** | Rv0659c-0660c (MazEF2) | **57** | Rv2546-2545 (VapBC18) |
| **18** | Rv0661c-0662c (VapBC7) | **58** | Rv2548-2547 (VapBC19) |
| **19** | Rv0665-0664 (VapBC8) | **59** | Rv2549c-2550c (VapBC20) |
| **20** | Rv0749-0748 (VapBC31) | **60** | Rv2596-2595 (VapBC40) |
| **21** | Rv0836c-0837c* | **61** | Rv2602-2601A (VapBC41) |
| **22** | Rv0910-0909 | **62** | Rv2653c-2654c |
| **23** | Rv0919-0918 | **63** | Rv2757c-2758c (VapBC21) |
| **24** | Rv0960-0959A (VapBC9) | **64** | Rv2759c-2760c (VapBC42) |
| **25** | Rv1045-1044* | **65** | Rv2801c-2801A (MazEF9) |
| **26** | Rv1102c-1103c (MazEF3) | **66** | Rv2826c-2827c* |
| **27** | Rv1114-1113 (VapBC32) | **67** | Rv2829c-2830c (VapBC22) |
| **28** | Rv1242-1241 (VapBC33) | **68** | Rv2863-2862A (VapBC23) |
| **29** | Rv1246c-1247c (RelBE1) | **69** | Rv2866-2865 (RelBE2) |
| **30** | Rv1397c-1398c (VapBC10) | **70** | Rv2872-2871 (VapBC43) |
| **31** | Rv1495-1494 (MazEF4) | **71** | Rv3180c-3181c (VapBC45) |
| **32** | Rv1546-1545 | **72** | Rv3182-3183 (HigBA3) |
| **33** | Rv1561-1560 (VapBC11) | **73** | Rv3189-3188 |
| **34** | Rv1720c-1721c (VapBC12) | **74** | Rv3320c-3321c (VapBC44) |
| **35** | Rv1741-1740 (VapBC34) | **75** | Rv3358-3357 (YefM/YoeB) |
| **36** | Rv1838c-1839c (VapBC13) | **76** | Rv3384c-3385c (VapBC46) |
| **37** | Rv1942c-1943c (MazEF5) | **77** | Rv3408-3407 (VapBC47) |
| **38** | Rv1953-1952 (VapBC14) | **78** | Rv3697c-3697A (VapBC48) |
| **39** | Rv1955-1956-1957 (TAC) | **79** | Rv3749c-3750c (VapBC50) |
| **40** | Rv1959c-1960c (ParDE1) |  |  |

### 2.3. Statistical analysis

The association of the polymorphisms in the TA systems to a particular drug resistance-based category was analysed by paired *t*-test with significance established at P values ≤ 0.01 using MS Excel (2010).

## 3. RESULTS

### 3.1. Polymorphisms in the TA genes and promoters

The polymorphisms within the toxin-antitoxin modules and the promoter regions were examined for 79 known and putative TA systems using H_37_Rv as a reference. Polymorphisms were detected in 80% (63/79) TA systems. The polymorphisms were primarily observed at up to 2 loci in a majority of the TA systems; however, few TA systems had polymorphisms at more than 2 loci. A maximum of 12 loci polymorphisms were observed in Rv0960-0959A (VapBC9), followed by 8 loci polymorphisms in Rv0059-0060 and Rv0836c-0837c. Overall, a comparatively higher number of polymorphic loci were localised in the toxin genes (58.7%) as compared to the antitoxin genes (40.7%). A single isolate exhibited polymorphism in the gene encoding the chaperone.

Single nucleotide polymorphism (SNP) reflecting transition mutation was found in 67.3% of the polymorphic loci. The second major type of polymorphism demonstrated transversion effect; while substitution, insertion and deletion were associated with a limited number of loci. Non-synonymous substitutions were observed in about 66% of the polymorphic sites in the TA genes; other polymorphisms resulted in either truncation or frameshift mutations. Truncation was indicated in the antitoxin genes of TA systems, Rv0059-0060 and Rv1495-1494 (MazEF4).

Of the 74 isolates analysed, 97.3% (72/74) isolates had at least one polymorphism in one of the TA systems. About 95% of the isolates exhibited a SNP R21R in RelBE2 TA system followed by Rv0919-0918 F141F (94.6%), VapBC47 S46L (94.6%) and VapBC22 A56V (90.5%) with a polymorphism at a particular locus which maybe annotated as a hotspot. In case of the promoters, 28 isolates exhibited polymorphism in the VapBC18 promoter sequence. Details of the number and type of polymorphisms, and the resultant change in the codon are summarised in Tables 2 and 3.

**Table 2.**
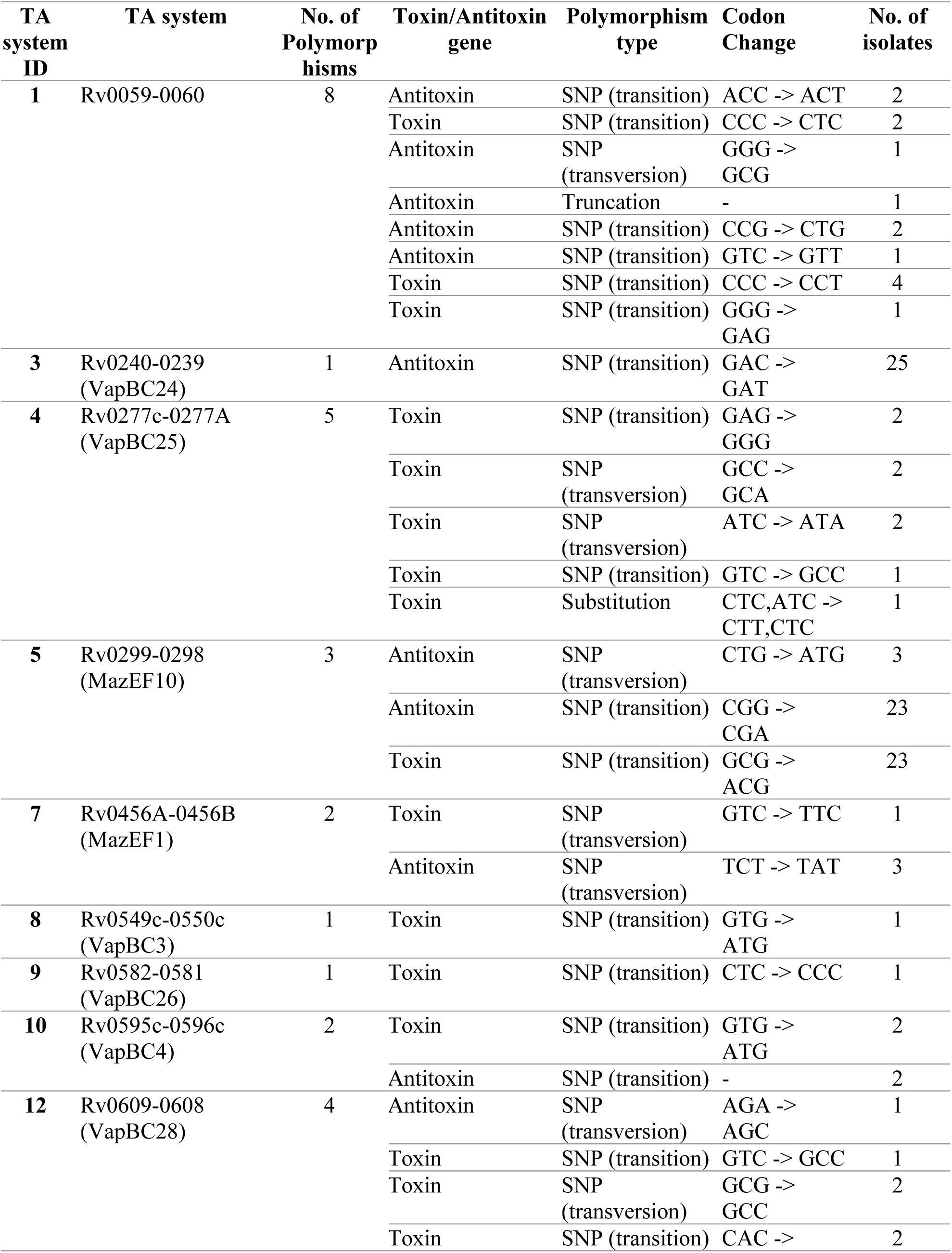
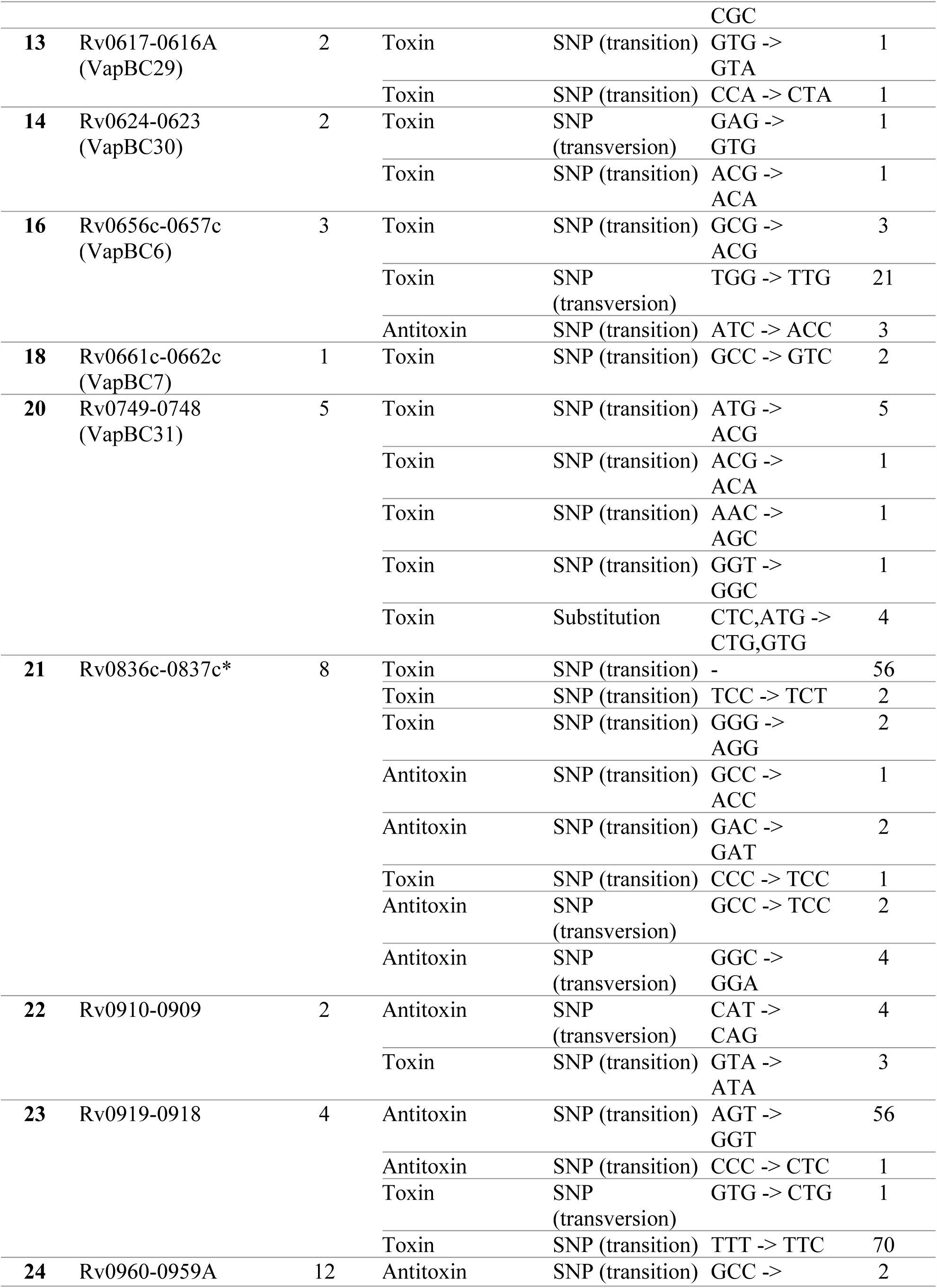
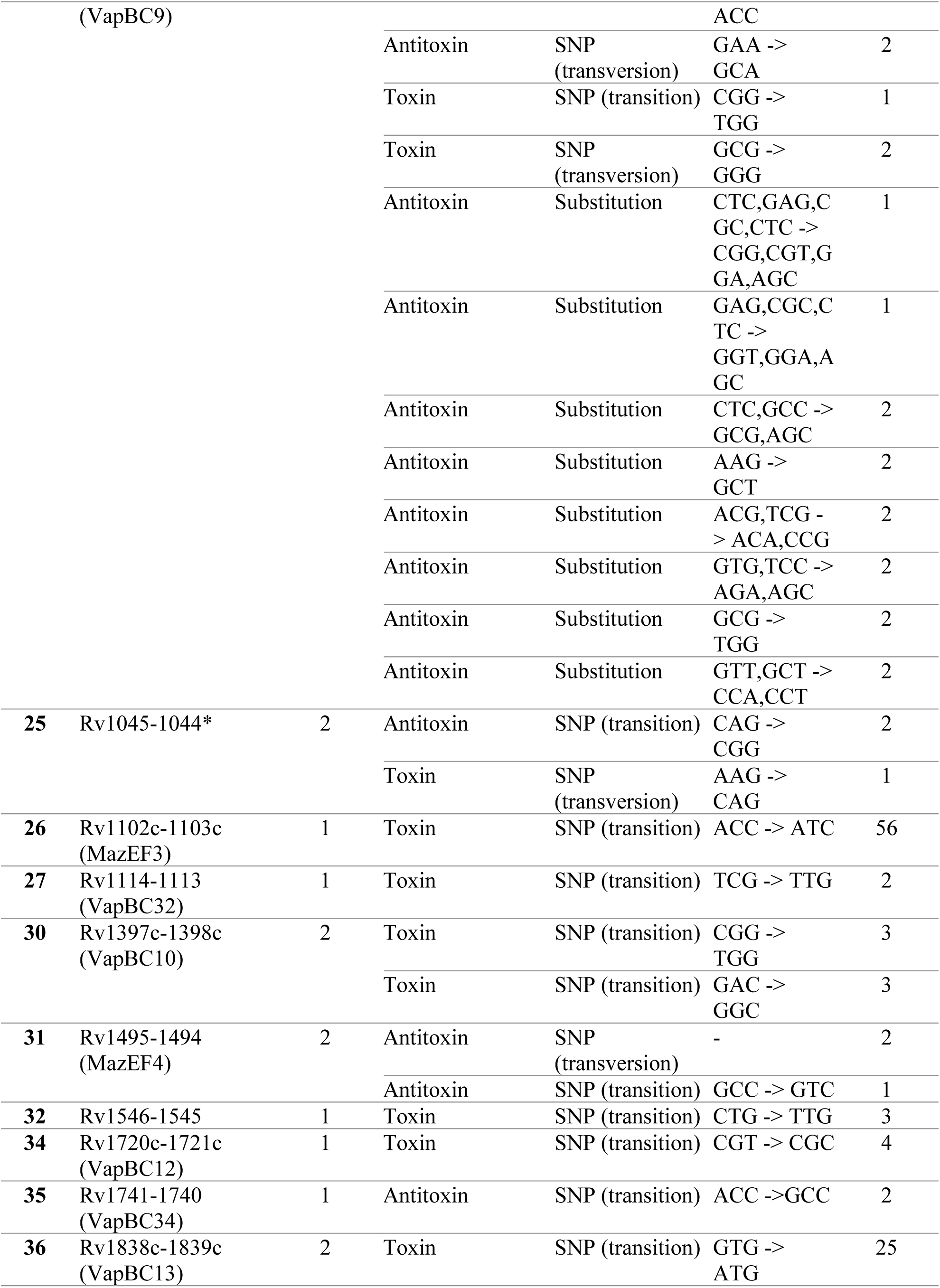
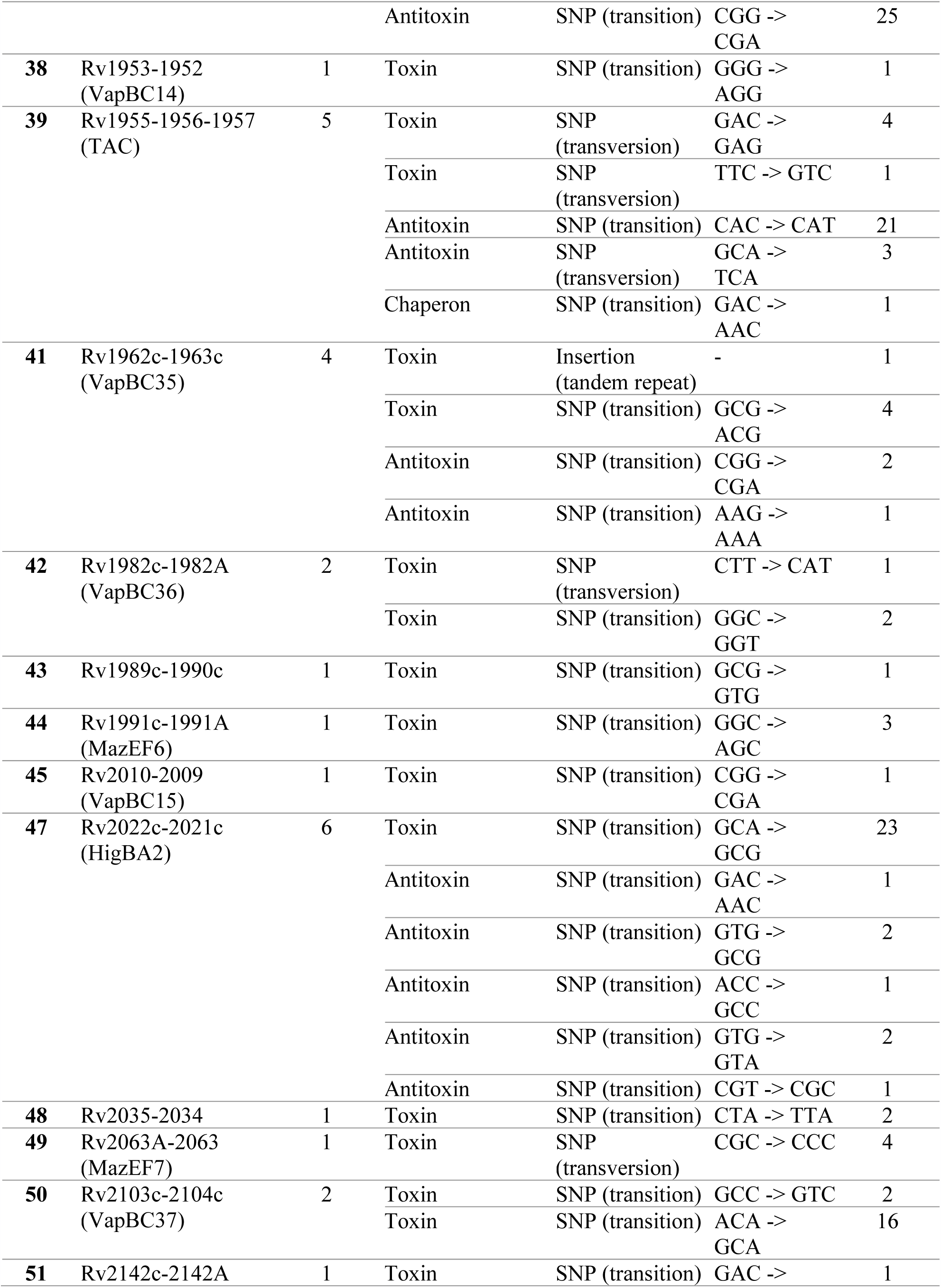
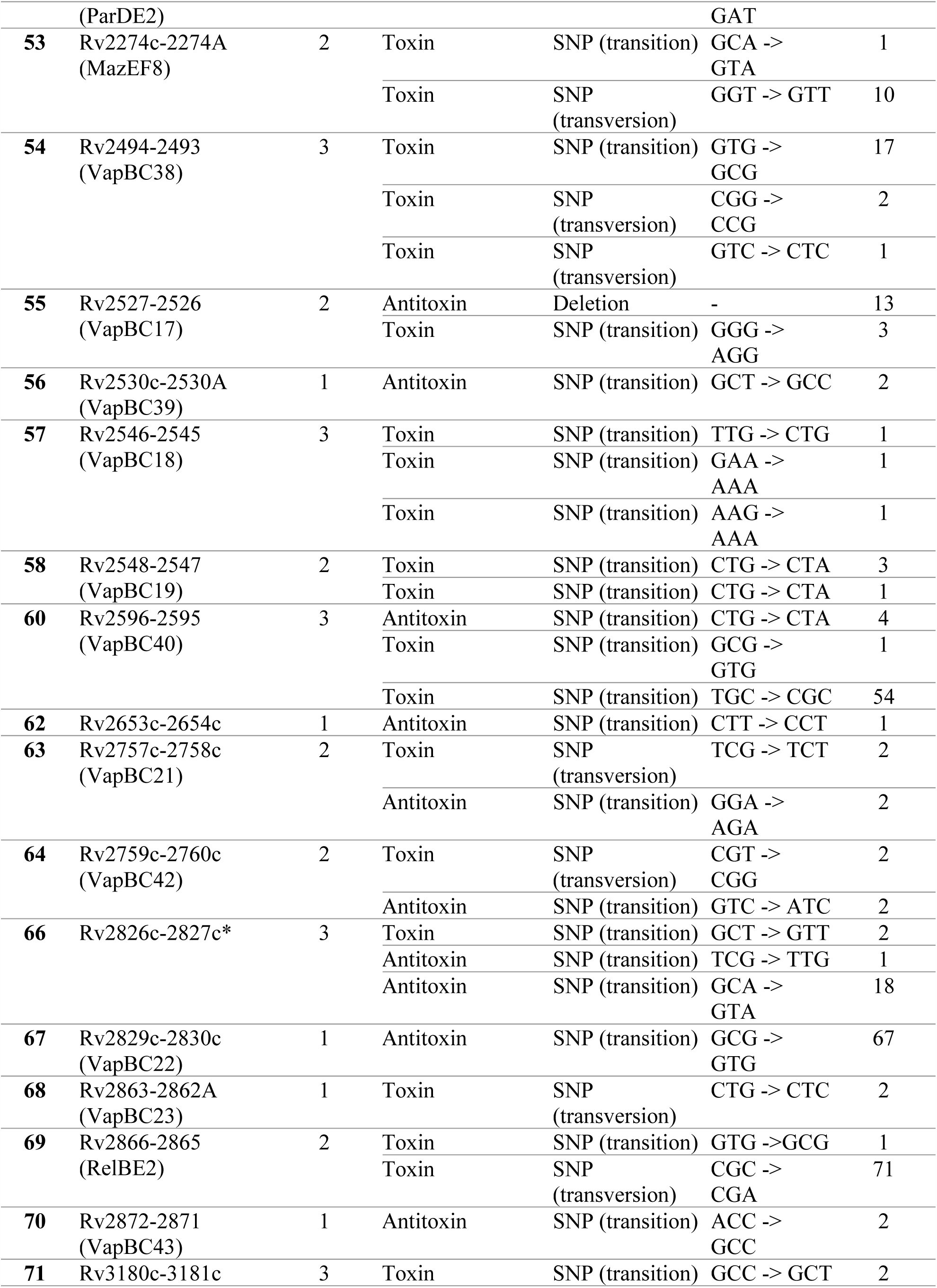
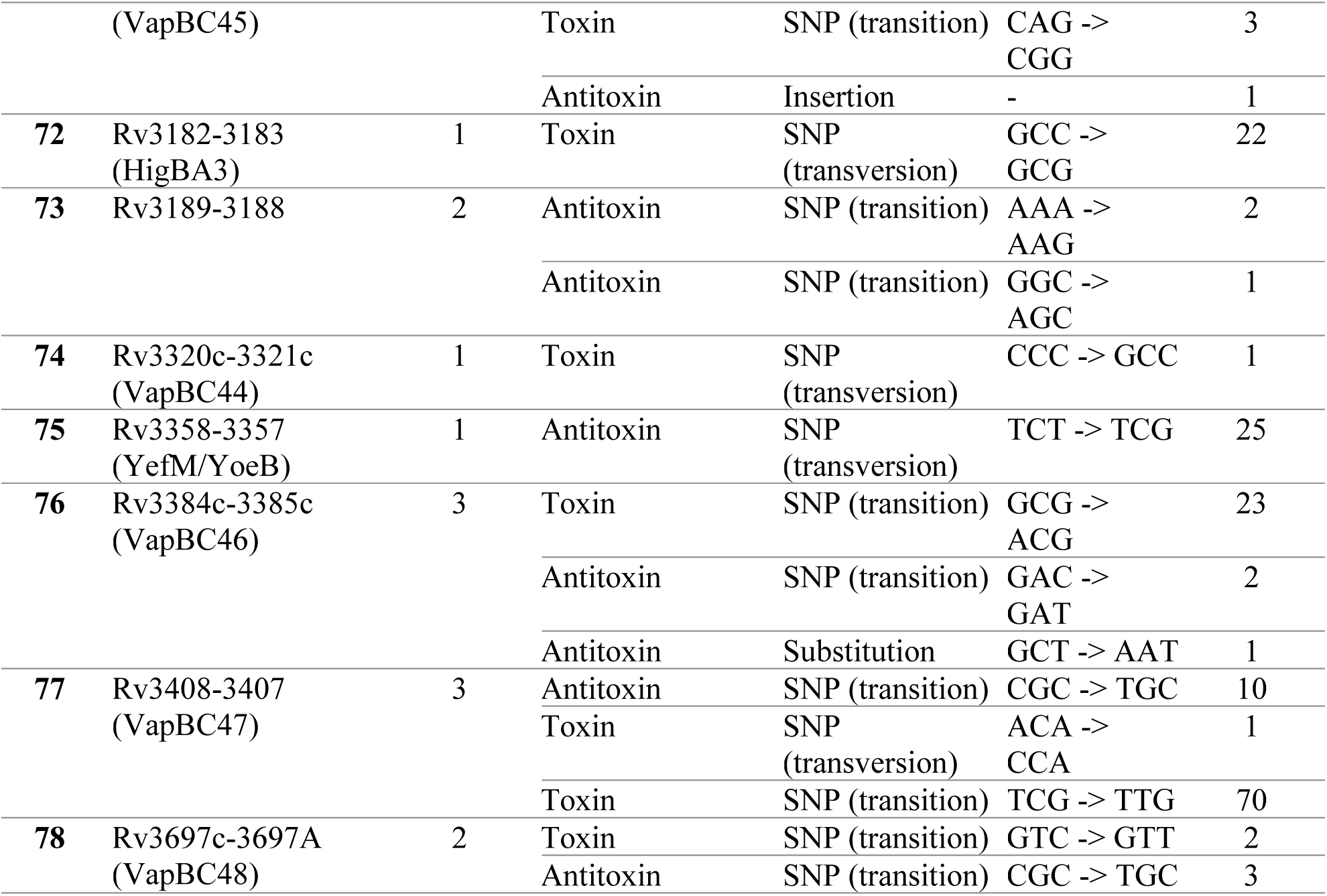
Polymorphisms in the TA genes.

**Table 3.** Polymorphisms in the promoters of the TA systems.

| TA system ID | TA system | No. of unique variant | Types of polymorphisms | Codon Change | No. of isolates |
| --- | --- | --- | --- | --- | --- |
| 2 | Rv0065-0064A (VapBC1) | 1 | Substitution | GGT→ AGT | 2 |
| 20 | Rv0749-0748 (VapBC31) | 1 | None | GGC → GGT | 7 |
| 24 | Rv0960-0959A (VapBC9) | 1 | Substitution | CTG → ATG | 1 |
| 57 | Rv2546-2545 (VapBC18) | 4 | None | GAC → GAT | 1 |
|  |  |  | None | CCC → CCG | 1 |
|  |  |  | None | ATT → ATC | 25 |
|  |  |  | None | CCG → CCA | 1 |
| 63 | Rv2757c-2758c<br>(VapBC21) | 1 | None | CGT -> CGG | 2 |

### 3.2. Polymorphisms in TA systems and drug susceptibility pattern

On the basis of the drug susceptibility data obtained using *Mykrobe Predictor*, the isolates were classified into Drug sensitive (DS) and Drug Resistant (DR) strains including multidrug resistant TB (MDR, resistant to isoniazid and rifampicin), pre-extensively drug resistant TB (pre-XDR, resistant to quinolones or any of the second line injectable in addition to isoniazid and rifampicin) and extensively drug resistant TB (XDR, resistant to quinolone and second line injectable along with isoniazid and rifampicin). The analysis for the presence of TA system polymorphisms in the genes of the strains with different categories using paired t-test revealed significant correlation between DS and DR strains detailed in Table 4. No significant difference was found within the DS and DR strains with respect to polymorphisms in the promoter sequences of the TA systems.

**Table 4.** Polymorphisms associated with DS and DR strains.

| TA system<br>ID | TA system | Polymorphisms | Association | p value |
| --- | --- | --- | --- | --- |
| 3 | Rv0240-0239<br>(VapBC24) | *vapB*24 D50D | DS | 0.0056 |
| 36 | Rv1838c-1839c<br>(VapBC13) | *vapC*13 V1M/<br>*vapB*13 R87R | DS | 0.0056 |
| 41 | Rv1962c-1963c<br>(VapBC35) | *vapC*35 A109T/<br>*vapC*35 Ins133/<br>*vapB*35 R89R/<br>*vapB*35 K33K | DR | 0.0059 |
| 75 | Rv3358-3357<br>(YefM/YoeB) | *yefM* S79S | DS | 0.0056 |

### 3.3. Polymorphisms in TA systems and strain lineages

A total of 27 isolates (36.5%) belonged to Beijing/East Asia lineage, followed by 25 (33.8%) Delhi/Central Asia, 17 (23%) European/American and 5 (6.8%) East Africa/Indian Ocean lineage. On analysing the presence of polymorphisms in various TA systems across the lineages, it was noted that specific polymorphisms in the TA systems were common in all the lineages; while others were specific to certain lineages. Polymorphisms in Rv0919-0918, Rv2829c-2830c (VapBC22), Rv2866-2865 (RelBE2) and Rv3408-3407 (VapBC47) were observed across all isolates irrespective of their lineage. The polymorphisms found exclusively or having significant association to specific lineages are presented in Table 5. With respect to the promoter sequences, polymorphisms in Rv2546-2545 (VapBC18) at a single locus was found to be associated to Beijing/East Asia lineage with 88.88% (n=24) prevalence rate. However, presence of additional polymorphisms at 3 other loci in the same sequence was found in one isolate which belonged to the East Africa/Indian Ocean lineage.

**Table 5.**
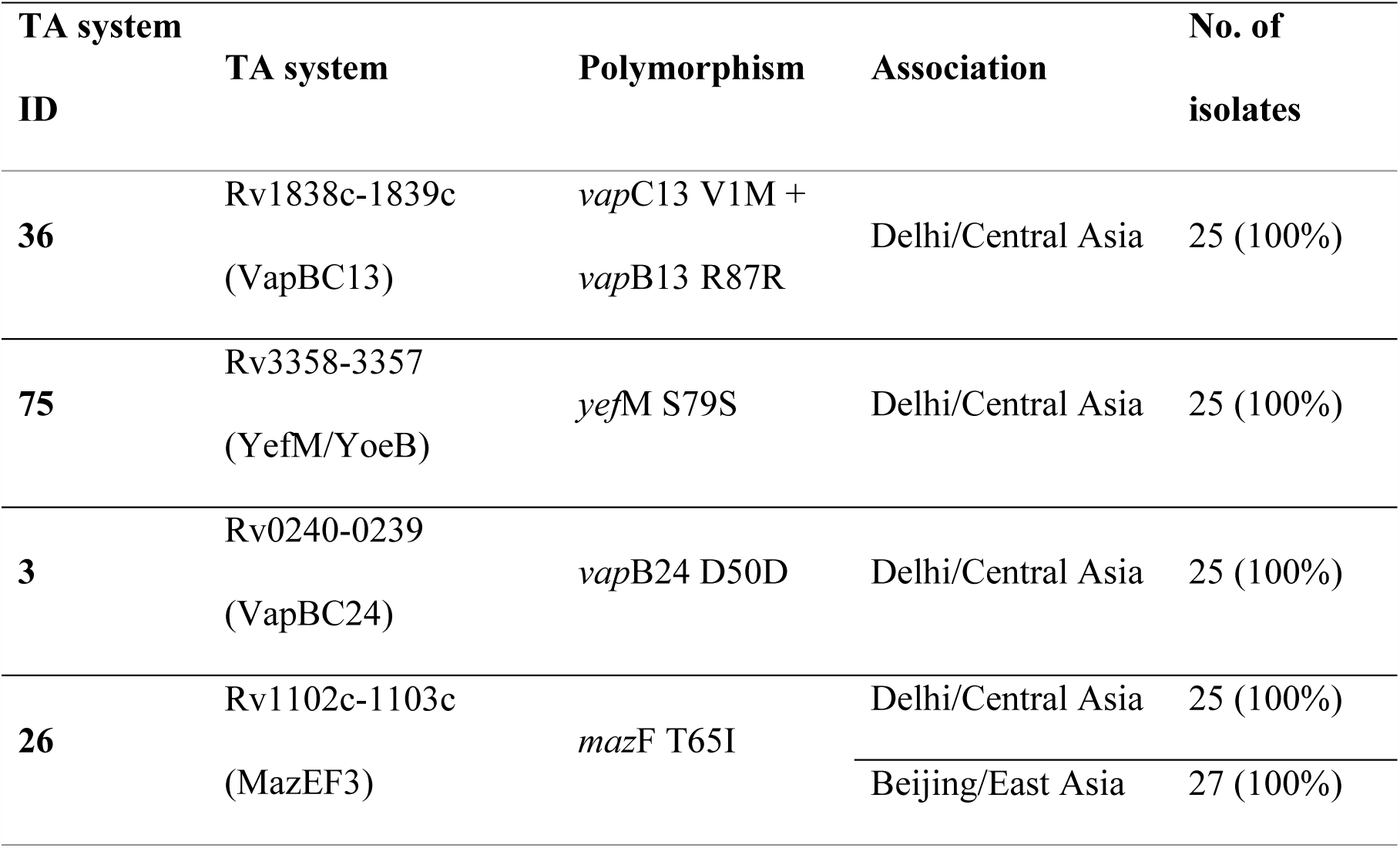
Polymorphisms associated with strain lineages.

| **TA system ID** | **TA system** | **Polymorphism** | **Association** | **No. of isolates** |
| --- | --- | --- | --- | --- |
| **36** | Rv1838c-1839c<br>(VapBC13) | *vapC13* V1M +<br>*vapB13* R87R | Delhi/Central Asia | 25 (100%) |
| **75** | Rv3358-3357<br>(YefM/YoeB) | *yefM* S79S | Delhi/Central Asia | 25 (100%) |
| **3** | Rv0240-0239<br>(VapBC24) | *vapB24* D50D | Delhi/Central Asia | 25 (100%) |
| **26** | Rv1102c-1103c<br>(MazEF3) | *mazF* T65I | Delhi/Central Asia | 25 (100%) |
|  |  |  | Beijing/East Asia | 27 (100%) |

Another interesting aspect in TA systems was polymorphism in the overlapping region between the toxin and the antitoxin genes. The overlapping region is found in 55/79 TA systems with regions varying from 1 bp to 17 bp. However, polymorphisms in the overlapping region were observed in 2 TA systems namely - Rv1838c-1839c (VapBC13) and Rv0595c-0596c (VapBC4), resulting in 2 different protein effects. In case of VapBC13, a substitution in the toxin VapC13 leading to a change in valine to methionine was observed; while in the antitoxin gene *vapB13*, no effect on the protein was observed. Similarly, in Rv0595c-0596c (VapBC4), a substitution of valine with methionine in the toxin VapC4 and truncation in the *vapB4* antitoxin gene were observed.

## 4. DISCUSSION

The persistence is principally mediated by the toxin-antitoxin systems in the bacteria [11]. The TA systems play a prominent role in development of persister cells rather than drug resistant cells [12]. There is limited correlation between the polymorphism in TA systems and drug susceptibility. A significant association was observed in 4/79 TA systems. Paired t-test revealed significant association of the presence of polymorphisms within Rv1962c-1963c (VapBC35) with respect to DR strains. An earlier study has shown altered expression pattern of VapBC35 TA module on exposure to antibiotics; this may be correlated to the polymorphisms associated with the DR strains [13]. Also, presence of polymorphisms in Rv0240-0239 (VapBC24), Rv1838c-1839c (VapBC13) and Rv3358-3357 (YefM/YoeB) are significantly associated with the DS strains of *M. tuberculosis*.

Polymorphisms resulting in substitution mutations observed within the overlapping regions of Rv1838c-1839c (VapBC13) and Rv0595c-0596c (VapBC4) resulted in a change from valine to methionine (GTG to ATG). Based on codon usage in *M. tuberculosis*, the codon change of GTG to ATG would likely result in higher translation rates of the toxins VapB13 and VapB3 [14]. Association of polymorphisms in the TA systems with the strain lineages was observed in case of the TA systems. In a novel finding in the present study, we showed association of polymorphism in Rv1838c-1839c (VapBC13), Rv3358-3357 (YefM/YoeB) and Rv0240-0239 (VapBC24) to Delhi Central Asia strains. The unique polymorphisms related to each of the three TA systems were observed exclusively in 100% (25/25) of the Delhi Central Asia strains. The polymorphic loci of all the 3 TA systems is localised in the antitoxin gene, with resultant silent (synonymous) mutations. Interestingly, polymorphisms in these TA systems are significantly associated with drug sensitive strains. Also, a unique SNP in Rv1102c-1103c (MazEF3) was observed in all the Beijing and Delhi Central Asia strains. A previous study has linked the SNPs in Toxin-Antitoxin-Chaperon (TAC) Rv1955-1956-1957 system with Beijing strains [15]. In the present study, 77% (21/27) Beijing strains exhibited polymorphism in the TAC system. With respect to the promoter sequences, polymorphism in Rv2546-2545 (VapBC18) at a single locus associated Beijing/East Asia lineage with a high (88.9%; 24/27) prevalence rate. Therefore, SNPs in TA systems may serve as targets in identification of the lineage of *M. tuberculosis* as reported by earlier studies [2, 16].

An interesting aspect in TA systems, is polymorphism in the overlapping region between the toxin and the antitoxin genes. The overlapping region is found in 55/79 TA systems with regions varying from 1 bp to 17 bp. However, polymorphisms in the overlapping region was observed only in 2 TA systems namely - Rv1838c-1839c (VapBC13) and Rv0595c-0596c (VapBC4) resulting in 2 different protein effects. A future perspective is to analyze the effect of SNP on the stability of the toxin and antitoxin protein molecules, and to determine the promoter strength for the variants using various bioinformatics tools. Experimental analyses of the effect of the polymorphisms in the expression of the toxin and antitoxin molecules will provide a better perspective on the precise effect of the polymorphisms.

Artificial activation of the TA systems in general have been reviewed earlier in the direction of discovering toxin activators or peptide disruptors of toxin-antitoxin for their possible application in antibacterial strategy [17, 18]. The TA systems in *M. tb* have been reviewed for the multiple toxin-antitoxin systems for their mechanisms and potential role in physiology and virulence [6]. There is a dearth of studies towards testing the TA system(s) in *M. tb* as a potential therapeutic tool, due to the risk of transformation into persister bacteria. With the increase in the incidence of drug resistance and with limited tools to deal with such resistant strains, it may be worth looking at the TA systems to seek a solution. Such studies will be beneficial in selection of TA systems for potential application in therapeutics.

## ETHICS STATEMENT

Ethical clearance is not required for this study.

## ACKNOWLEDGMENTS

The authors would like to thank Dr. Nerges Mistry, The Foundation for Medical Research (FMR) and Dr. Dhananjaya Saranath, Consultant, FMR, for critical reading of the manuscript.

